# Carbon nanofibers fabrication, surface modifications, and application as the innovative substrate for electrical stimulation of neural cell differentiation

**DOI:** 10.1101/2022.10.15.512333

**Authors:** Houra Nekounam, Hadi Samadian, Hossein Golmohammadi, Fatemeh Asghari, Mohammad Ali Shokrgozar, Samad Ahadian, Reza Faridi Majidi

**Affiliations:** Department of Medical Nanotechnology, School of Advanced Technologies in Medicine, Tehran University of Medical Sciences, Tehran, Iran; Department of Molecular Medicine, School of Medicine, Hamedan University of Medical Sciences, Hamedan, Iran; Department of Electrical and Computer Engineering, Islamic Azad University of Tehran-North, Tehran, Iran; National Cell Bank of Iran, Pasteur Institute of Iran, Tehran, Iran; NouBio Inc., Los Angeles, USA

**Keywords:** CNFs, Surface functionality, Neural tissue engineering, Electrical stimulation

## Abstract

Engineered nanostructures are innovative and precisely designed, synthesized, and tailored with outstanding physicochemical properties that can be applied as the game-changer in neural tissue engineering. The present study aimed to develop an innovative approach based on electrical stimulation through a conductive scaffold to differentiate neural cells from human adipose mesenchymal stem cells without the use of a specific environment for neural differentiation. Electrospun carbon nanofibers (CNFs) were obtained using heat treatment of polyacrylonitrile nanofibers and treated by nitric acid, ethylenediamine, and oxygen Plasma. SEM imaging revealed that the treated nanofibers have s diameter in the range of 120-200 nm and the treatment did not significantly change the CNFs diameter. The FTIR results showed that the treatments were able to introduce COOH, OH, and NH2 functional groups on the CNFS surface. The XRD and Raman analysis showed that the plasma treatment induced the lowest structural changes in the CNFs microstructure. The biocompatibility assessments showed that the pristine and treated CNFs were non-toxic induced proliferative effect on human adipose-derived mesenchymal stem cells. The electrical stimulation (1.5 mA current with a frequency of 500 Hz and CMOS waveform for 7 days 10 min each day) induced the expression of neural genes and proteins by the cells cultured on the treated CNFs. The Plasma-treated CNFs mediated the highest differentiation outcome. These results indicate that electrospun CNFs can be applied as the innovative interface applicable for neural tissue regeneration under electrical stimulation.

**Research highlights:**

- CNFs were fabricated from PAN nanofibers
- Different amounts of ZnONPs were incorporated into or sprayed on CNF
- increasing in ZnONP amount decreased conductivity, surface wettability was improved by ∼19–33%.
- Also, FTIR, XRD, and Raman analyses proved that the presence of ZnONP improved structure formation with lower defect density

Schematic 1.
The applied electrical stimulation setup

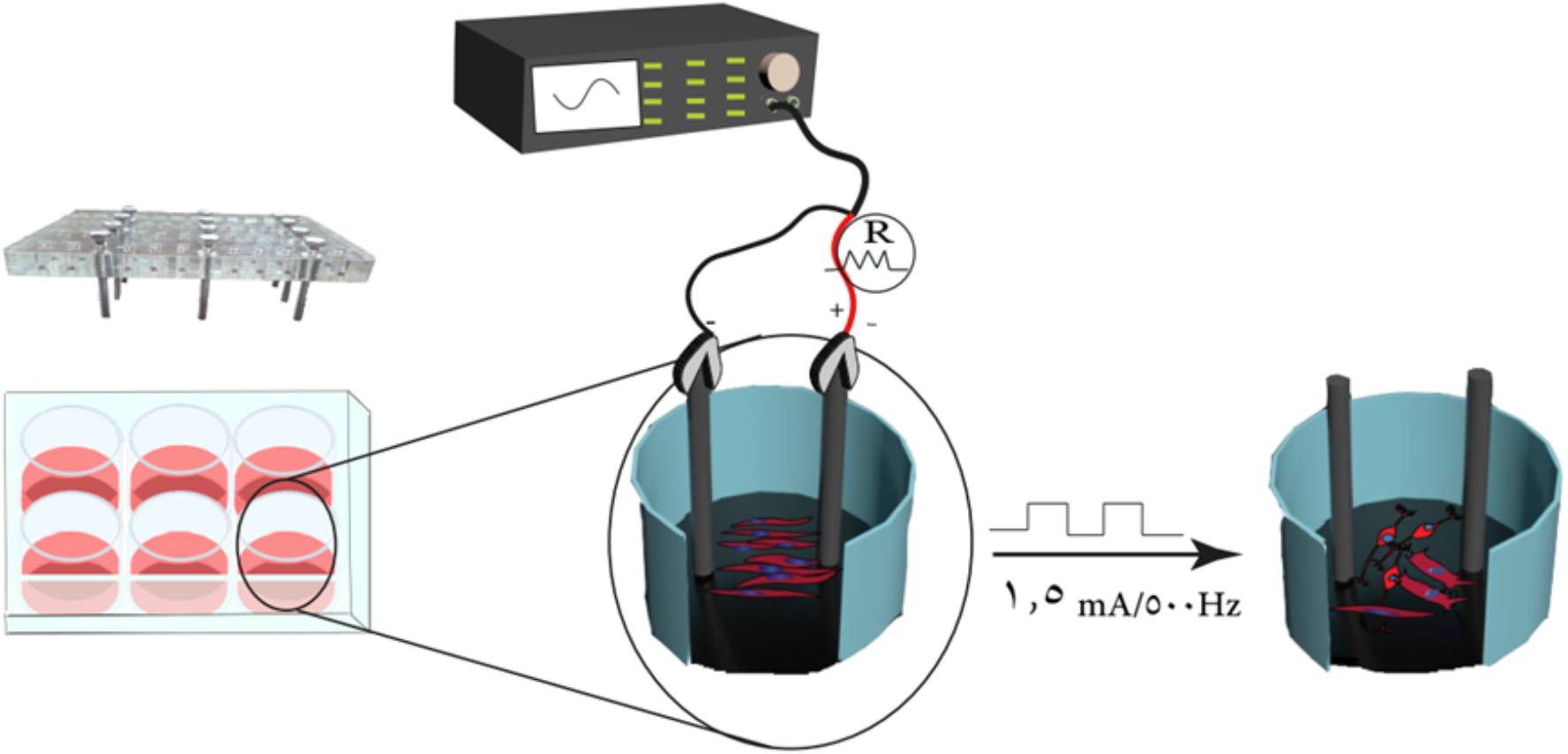

## 1. Introduction

Injuries to the nervous systems (both the central nervous system (CNS) and peripheral nervous system (PNS)) inducing temporal or permanent disabilities have substantial negative effects on the population and have a huge burden on the medical and healthcare system, as well as the social and economic field (1, 2). The frequency of such damages has been growing due to modern lifestyles. Moreover, in mammals, the autonomous regeneration capacity of both CNS and PNS is low and life-lasting disabilities are common after disease and injury. Accordingly, conventional treatments are not completely successful and effective. Moreover, neural tissue regeneration is highly complicated and dependent on a variety of cellular, structural, and biochemical factors. Therefore, sophisticated and complementary treatment approaches are required to betaine the highest anatomical and functional recovery. The implantation of autologous nerve grafts is the gold standard for PNS injuries. Despite positive treatment outcomes, this approach suffers from several issues such as loss of function at the donor site, donor site pain, limited donor tissue, an increase of operative cost and time, and mismatch of the damaged nerve and graft size (3-5). Therefore, innovative approaches have been proposed as alternative treatment options to provide better treatment outcomes and eliminate the shortcomings of conventional treatments.

It has been reported that electrical stimulation (ES) has promising effects on neural cells growth, differentiation, and functions (6, 7). The neural ES is based on the fact that endogenous bioelectricity presents in the human body has critical functions in maintaining normal biological functions of neural cells (8, 9). Moreover, the application of external has shown promising results in the clinic. For instance, deep brain stimulation is prescribed for patients suffering from movement disorders that no longer respond properly to medications (10). It requires the implantation of an array of electrodes in the brain that has complications and difficulties. Alternatively, using the electroconductive scaffolds, in the concept of tissue engineering, is an innovative approach with fascinating advantages.

The tissue engineering approach is the combination of biology, material sciences, chemical and physical cues, with engineering sciences/technologies (11, 12). The combination of ES with tissue engineering requires the electroconductive structures as the scaffolds and/or substrate to stimulate the cells attached on and/or infiltrated into the scaffolds (13). Nanotechnology as an enabling point of view has emerged to improve and correct the limitations of conventional strategies (14). The human body systems are working as the nanoscale and can be considered as the nanomachines, accordingly nanostructured scaffolds can interact more effectively with the cells and progress involved in the regeneration of damaged neural tissue (15, 16). Furthermore, it has been observed that nanostructured scaffolds are more potent than their bulk structure counterparts. Various types of electroconductive substances have been applied to fabricate scaffolds, such as metals, polymers, and carbon-based materials.

Electrospun carbon nanofibers (ECNFs) are one-dimensional structures with various promising properties beneficial for tissue engineerings, such as morphological resemblance to the extract cellular matrix (ECM) of tissues, fascinating mechanical and electrical properties, biocompatibility, and chemical inertness (17, 18). These features have made ECNFs promising candidates to be applied as the electroconductive scaffold for ES of neural cells. Despite their promising properties, CNFs suffer from surface hydrophobic and surface functionality (19). Accordingly, in the current study, we applied different surface modifications to tailor the chemical and physical properties of ECNFs beneficial for electrical stimulation of mesenchymal stem cells (MSCs) to differentiate toward neural cells.

## 2. Results and discussion

### 2.1. CNFs fabrication and characterization

The morphology of the CNFs was visualized using SEM and the results are presented in Figure 1. The results showed that the surface treatments did not induce any adverse effects on the morphology of the CNFs. Moreover, pristine CNFs have a diameter of 154± 30.5 nm and CNF/CCOH, CNF/NH2, CNF/Plasma, and CNF/OH have a diameter of 146.6 ± 35.0, 186.6 ± 75.4, 157.5 ± 28.6, and 153.3 ± 35.1 nm, respectively. It was observed that the treatments did not significantly change the diameter of CNFs. It can be concluded that the treatments affected just the surface of CNFs and not the overall structure of CNFs.

**Figure 1.**
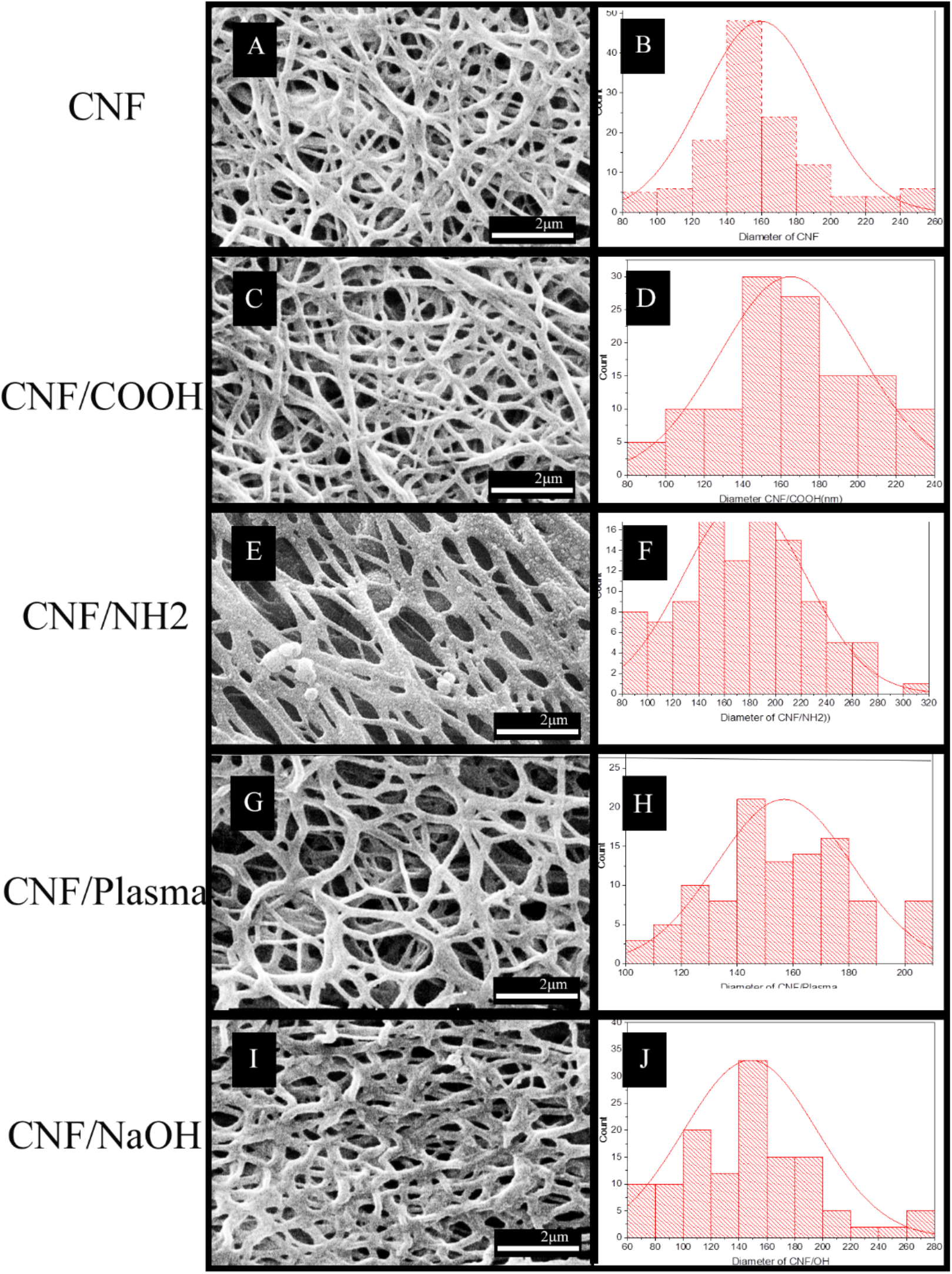
SEM micrograph of the CNFs. (A and B) pristine CNF; (C and D) CNF/CCOH; (E and F) CNF/NH_2_; (G and H) CNF/Plasma; (I and J) CNF/OH

The introduction of the intended surface functional groups and the chemistry of CNFs surface were evaluated using FTIR spectroscopy. The FTIR spectroscopy showed that the nitric acid treatment induced carboxylic groups on the CNFs (**Figure 2A**). The peaks located at 1000, 1083, 1301, and 1141 cm^-1^ are related to carboxylic groups and C-O vibrations. The peak located at 1727cm^-1^ is attributed to the aldehyde functional group (C=O). Moreover, acid treatment for 2 hr induced more OH dimer group of carboxylic acid (peaks at 2964, 3160, 3306, and 3380 cm^-1^) compared to 30 min and 1 hr acid treatments. It was observed that the longer treatment time destroyed the CNFs structures (data not shown).

**Figure 2.**
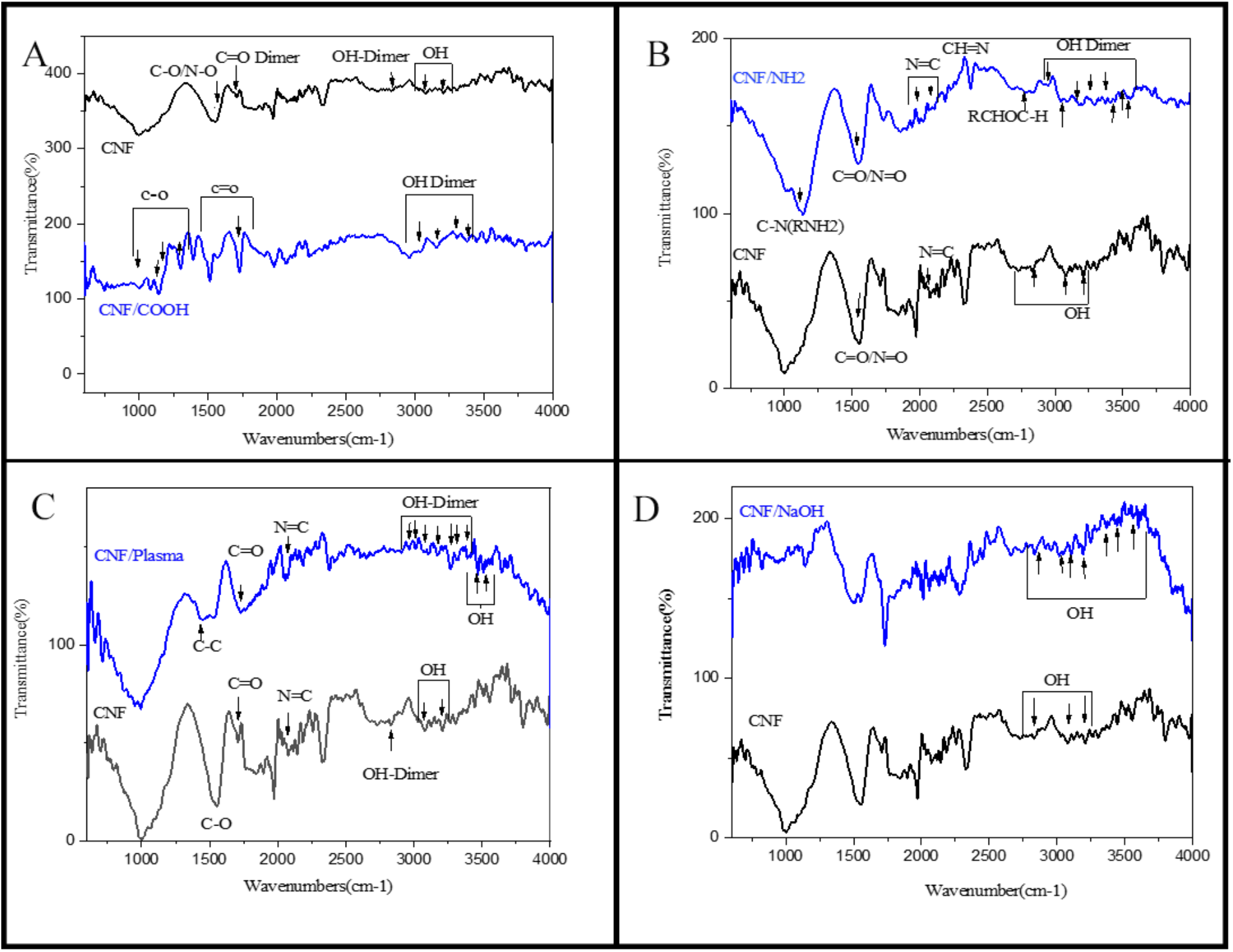
AT-FTIR spectrum and XRD patter of CNFs. (A) FTIR spectrum of carboxylation process at different time, (B) FTIR spectrum of amine-functionalized CNFs, (C) FTIR spectrum of plasma treated CNFs, and (D) FTIR spectrum of NaOH treated CNFs,

The carboxylated CNFs were then subjected to the amination process using EDA and the FRIT results confirmed the amination process. The sharp peak at 1136 cm^-1^ is related to NH2R or C-N groups and the peaks located at 1542, 1994, 2068, and 2229 cm^-1^ are indicating the N=O, R-N=C, N=C, and CH=N, respectively (**Figure 2B**). The oxygen Plasma treatment of CNFs induced plenty of hydroxyl functional groups on CNFs (indicated by the peaks at 2963, 3009, 3102, 3170, 3265, 3313, 3390, 3469, and 3508 cm^-1^) (**Figure 2C**).

The effect of the treatment on the crystallinity of the CNFs was evaluated using the XRD analysis and the results are presented in **Figure 3** (supplementary) and **Table 2**. The results showed that the peaks located at 26.5 (002 crystals plane) and 44.0 2Ө (001 crystals plane) indicate the presence of graphene nanocrystals in the structure of CNFs (20-22). Moreover, it was observed that the surface treatment reduced the crystallinity of CNFs. According to quantitative results presented in Table 1 and the full width at half maximum (FWHM) values extracted from XRD analysis, the NaOH and oxygen Plasma treatments induced the highest and lowest effects, respectively, on the crystallinity of CNFs which was stated in our previous study (23, 24).

**Table 1.** Quantitative crystallinity results

| Samples | 2Θ | FWHM | d(002) °A | Lc(nm) |
| --- | --- | --- | --- | --- |
| CNF/COOH | 23.83 | 0.29 | 3.73 | 3.6 |
| CNF/NH<sub>2</sub> | 26.54 | 0.39 | 3.35 | 2.4 |
| CNF/Plasma O<sub>2</sub> | 24.99 | 0.14 | 3.56 | 7.3 |
| CNF/OH | 28.31 | 0.59 | 3.15 | 1.4 |

**Table 2.** The quantitative structural results of CNFs derived from Raman analysis.

| Samples | D<br>Position<br>(cm<sup>-1</sup>) | G<br>Position<br>(cm<sup>-1</sup>) | ID/IG<br>(integrated<br>areas) | Crystallite<br>Parameters |  | Defect Parameters |  | R(ID/IG) |
| --- | --- | --- | --- | --- | --- | --- | --- | --- |
|  |  |  |  | Size<br>La(nm) | Area<br>La<sup>2</sup><br>(nm<sup>2</sup>) | Average<br>Distance<br>LD(nm) | Density<br>n<sub>D</sub>(cm-<br>2*10<sup>11</sup>) |  |
| CNF | 1372.14 | 1582.42 | 3.41 | 5.63 | 31.69 | 2.05 | 7.65 | 0.97 |
| CNF/NH2 | 1373.14 | 1592.75 | 2.77 | 6.93 | 48.02 | 2.28 | 6.21 | 1.08 |
| CNF/COOH | 1377.47 | 1587.59 | 2.83 | 6.79 | 46.10 | 2.25 | 6.35 | 1.01 |
| CNF/Plasma<br>o2 | 1366.78 | 1582.42 | 3.47 | 5.53 | 30.58 | 2.03 | 7.79 | 0.97 |
| CNF/OH | 1377.47 | 1587.59 | 3.62 | 5.30 | 28.09 | 1.99 | 8.26 | 1.04 |

**Figure 3.**
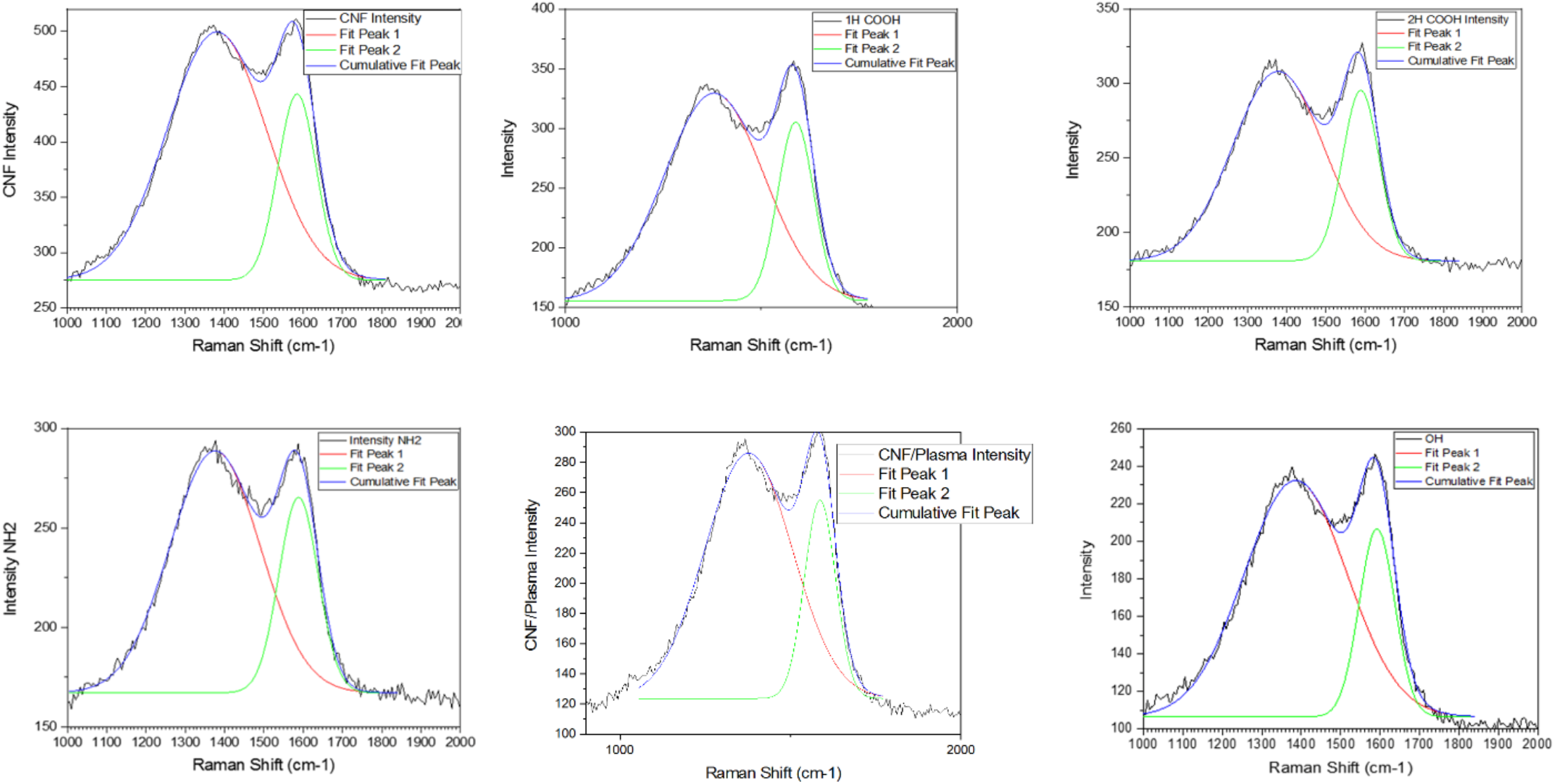
Raman spectra of CNFs with different surface functionalities.

Considering that in the plasma and sodium hydroxide methods, the hydroxide functional group is formed on the surface, the results confirm that the changes in the crystal structure are due to the selected method and not due to the functional group. On the other, since the difference between the carboxyl and amine group is low so most changes are due to the method of carboxylation.

The microstructure of the treated CNFs was investigated using Raman spectroscopy and the results are shown in **Figure 4**. and **Table 2**. The results showed that the treatments increased the graphitization of the CNFs due to the effects of the solvents on the amorphous graphitic regions on CNFs [26,27]. As shown in Figure 3, the surface treatments moved the D and G peaks to the higher wavenumbers. Moreover, increasing the D and G peaks intensity ratio reveals increasing the ratio of edges to crystal in and graphitic planes. The Plasma and NaOH treatments reduced the size of the crystals while increasing the density of defects. On the other hand, acid treatment and amination increased the size of the crystals while reducing the density of defects. It can be concluded that the solvent type and time of incubation have a direct correlation with the crystallinity and microstructure of CNFs and in this study, the oxygen Plasma treatment induced the lowest effect. In the method related to sodium hydroxide, the most structural degradation is observed and according to the Raman and XRD results, the plasma oxygen method is selected to create the hydroxide functional group.

**Figure 4.**
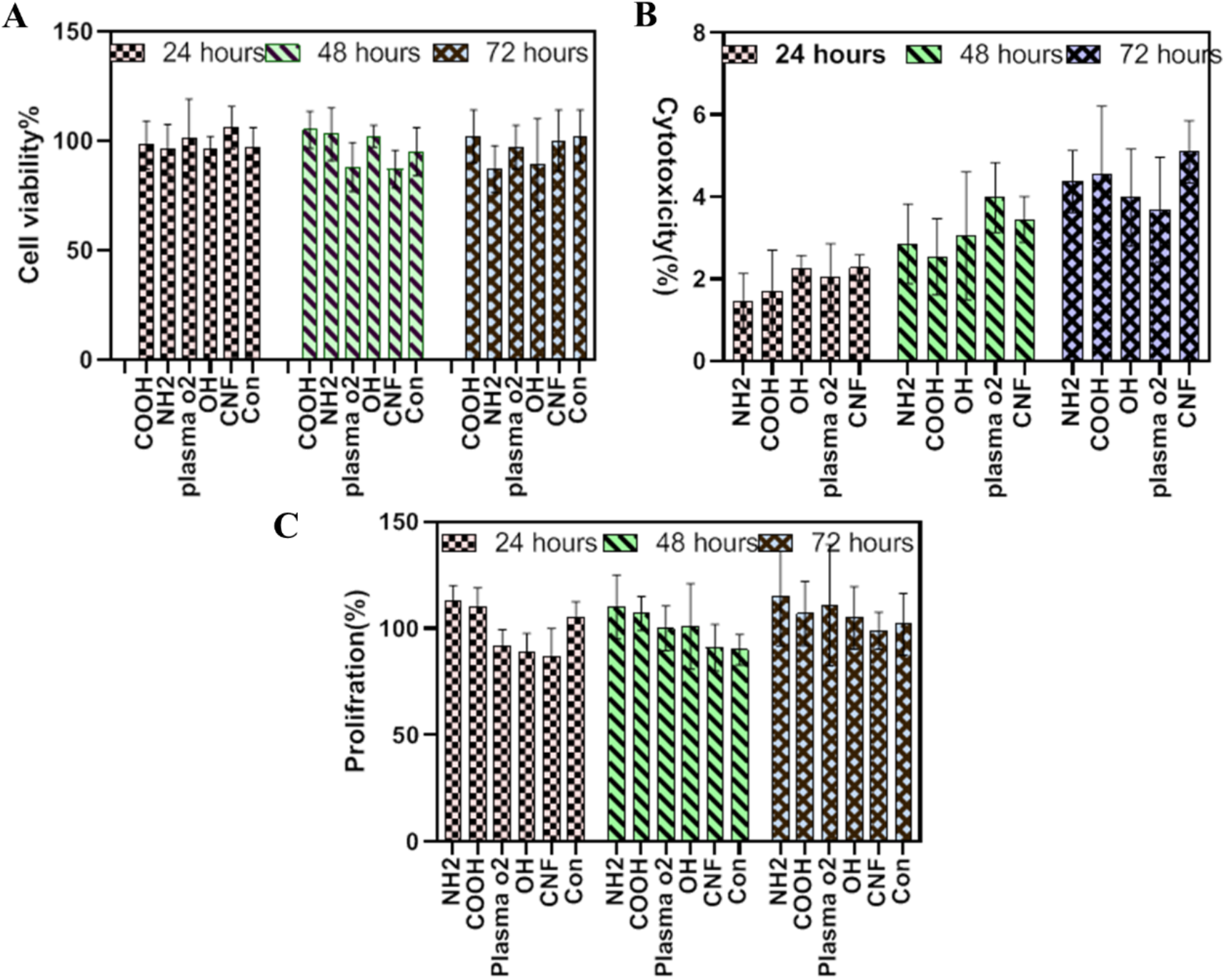
Viability/proliferation of hAMSC on CNFs different surface functionalities. (A) in-direct MTT result, (B) direct toxicity result based on LDH assay, and (D) proliferation results based on total LDH assay

The effects of the surface treatments on the electrical conductivity of the CNFs were investigated and observed that the treatments reduced the electrical conductivity of CNFs (**Table 3**). According to the FTIR, XRD, and Raman results, the electrical conductivity reduction can be related to the introduction of various surface functional groups on CNFs, as well as changes in the crystallinity.

**Table 3.**
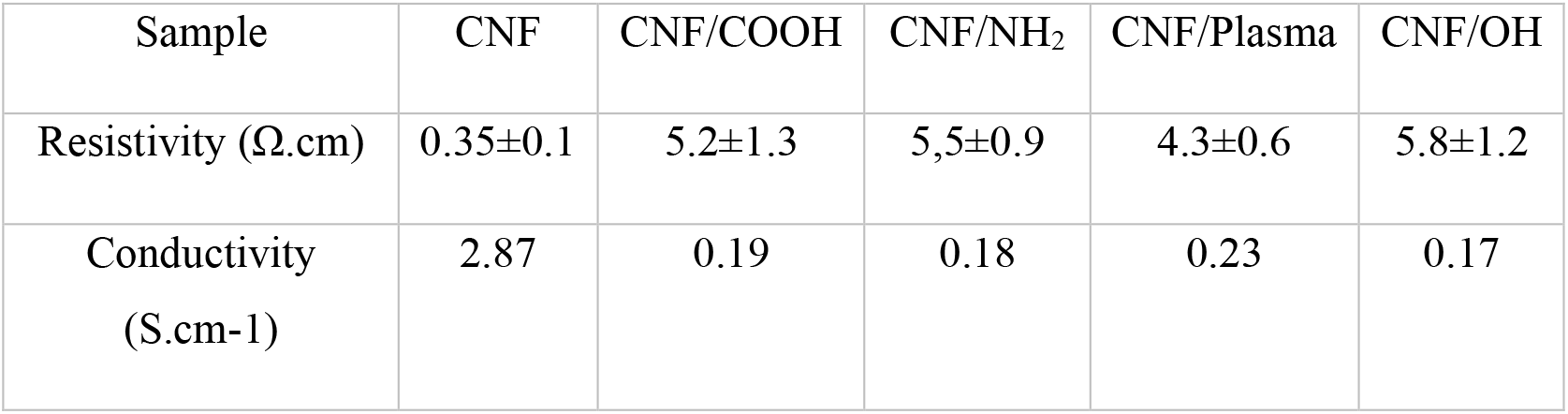
The electrical properties of the CNFs

| Sample | CNF | CNF/COOH | CNF/NH<sub>2</sub> | CNF/Plasma | CNF/OH |
| --- | --- | --- | --- | --- | --- |
| Resistivity ($\Omega\cdot\text{cm}$) | 0.35±0.1 | 5.2±1.3 | 5.5±0.9 | 4.3±0.6 | 5.8±1.2 |
| Conductivity<br>($\text{S}\cdot\text{cm}^{-1}$) | 2.87 | 0.19 | 0.18 | 0.23 | 0.17 |

The WCA measurement was conducted to assess the wettability of the CNFs and observed that the surface treatments reduced the WCA, indicating an improvement in surface hydrophilicity. As shown in Figure 5(Supplementary). The pristine CNFs surface is hydrophobic with a WCA value of 100 ± 9 °. The WCA value for CNF/COOH, CNF/NH2, CNF/Plasma, and CNF/OH was 72 ± 10, 32 ± 45, 12 ± 5, and 52 ± 27 °, respectively. These findings showed that the treatments improved the wettability of CNFs, as a critical parameter for biomedical applications.

**Figure 5.**
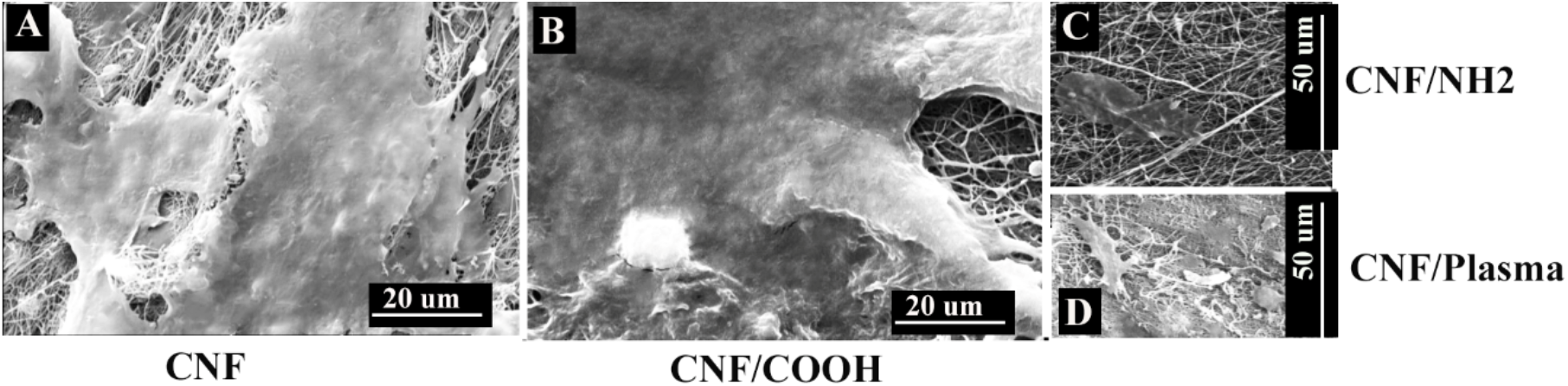
Morphology of hAMSCs on CNFs. (A) pristine CNFs, (B) CNFs-COOH, and (C) CNFs-NH2 (D) CNFs-Plasma O2

Mechanical properties of the CNFs were investigated based on the tensile strength method and the results are presented in **Figure 6**(supplemetary). and **Table 4**. The results showed that the CNFs were brittle and the surface treatments did not significantly alter the strain-stress curve of CNFs. According to the values stated in the previous study [26,27]. Although, the ultimate tensile strain (UTS) value of pristine CNFs and CNF/NH2 were in the same range but reduced for CNF/OH, CNF/Plasma, and CNF/COOH. It can be concluded that the, although the surface treatment modified the surface chemistry of CNFs, did not compromised the tensile strength of the CNFs.

**Table 4.** Mechanical properties parameters of CNFs derived from the Stress-Stain curves

| Samples | CNF | COOH | NH<sub>2</sub> | Plasma | OH |
| --- | --- | --- | --- | --- | --- |
| **Module (Mpa)** | 1338± 61 | 1429± 53 | 1338± 78 | 1364± 64 | 1417± 32 |
| **R2** | 0.99 | 0.99 | 0.99 | 0.99 | 0.99 |
| **UTS (MPa)** | 11.58± 5.2 | 5.96± 2.3 | 11.31± 4.3 | 2.76± 1.7 | 2.42± 0.98 |
| **Elongation at break (%)** | 3.16± 2.4 | 7.42± 1.4 | 3.00± 1 | 2.40± 1.5 | 4.94± 1.2 |

**Figure 6.**
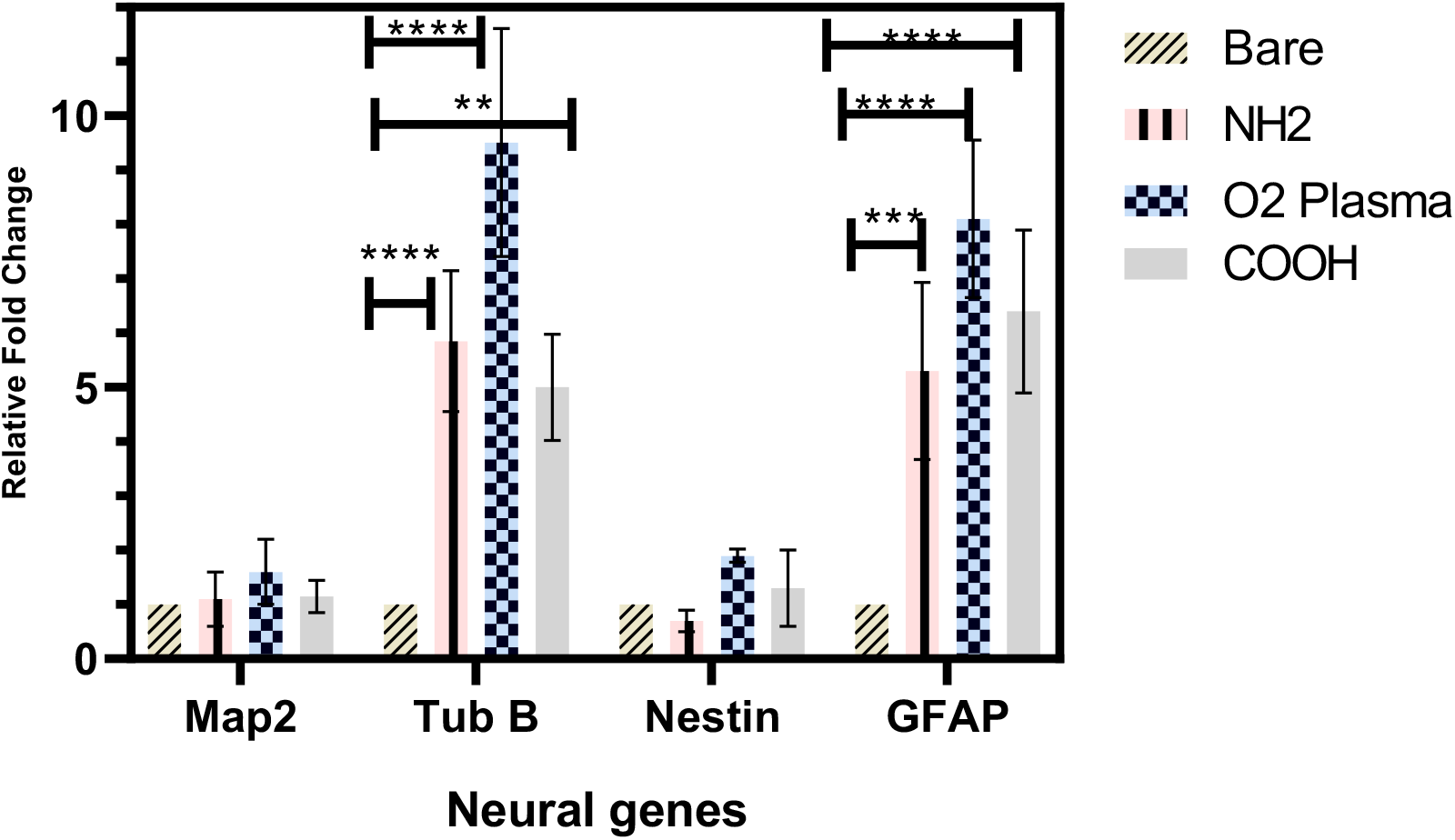
Expression of neural gens on CNFs with different surface functionalities.

### 2.2. Biological studies results

#### 2.2.1. Cell viability/proliferation on CNFs

The toxic effect of CNFs on the cells was evaluated using the MTT assay kit (in-direct approach) and the LDH activity assay kit (direct approach). The results of the in-direct MTT assay (**Figure. 7A**) showed that the treated CNFs did not leak any harmful substance to the extract and did not induce significant toxicity (p < 0.05). Although the toxicity of the treated CNFs was more than the pristine CNFs, that can be due to the induced structural defect during the treatments, which enhances substances leakage from the defects. The direct toxicity evaluation using the LDH activity assay kit showed that the induced toxicity was lower than 7% and the treated CNFs were more biocompatible than the pristine CNFs (**Figure 7B**). The proliferation of the cells on the CNFs evaluated by the total LDH activity assay approach (**Figure 7B)** showed that the treated CNFs induced a more proliferative effect which was not statistically significant (p < 0.1). These observations revealed that, although the treated CNFs induced a little more in-direct toxic effect, they support direct cell-scaffold attachment and induced a more proliferative effect, which can be related to the induction of various surface functional groups beneficial for cell attachment and growth. Moreover, enhanced surface hydrophilicity induced by the surface treatments improves the cell’s attachment and growth on the CNFs.

**Figure 7.**
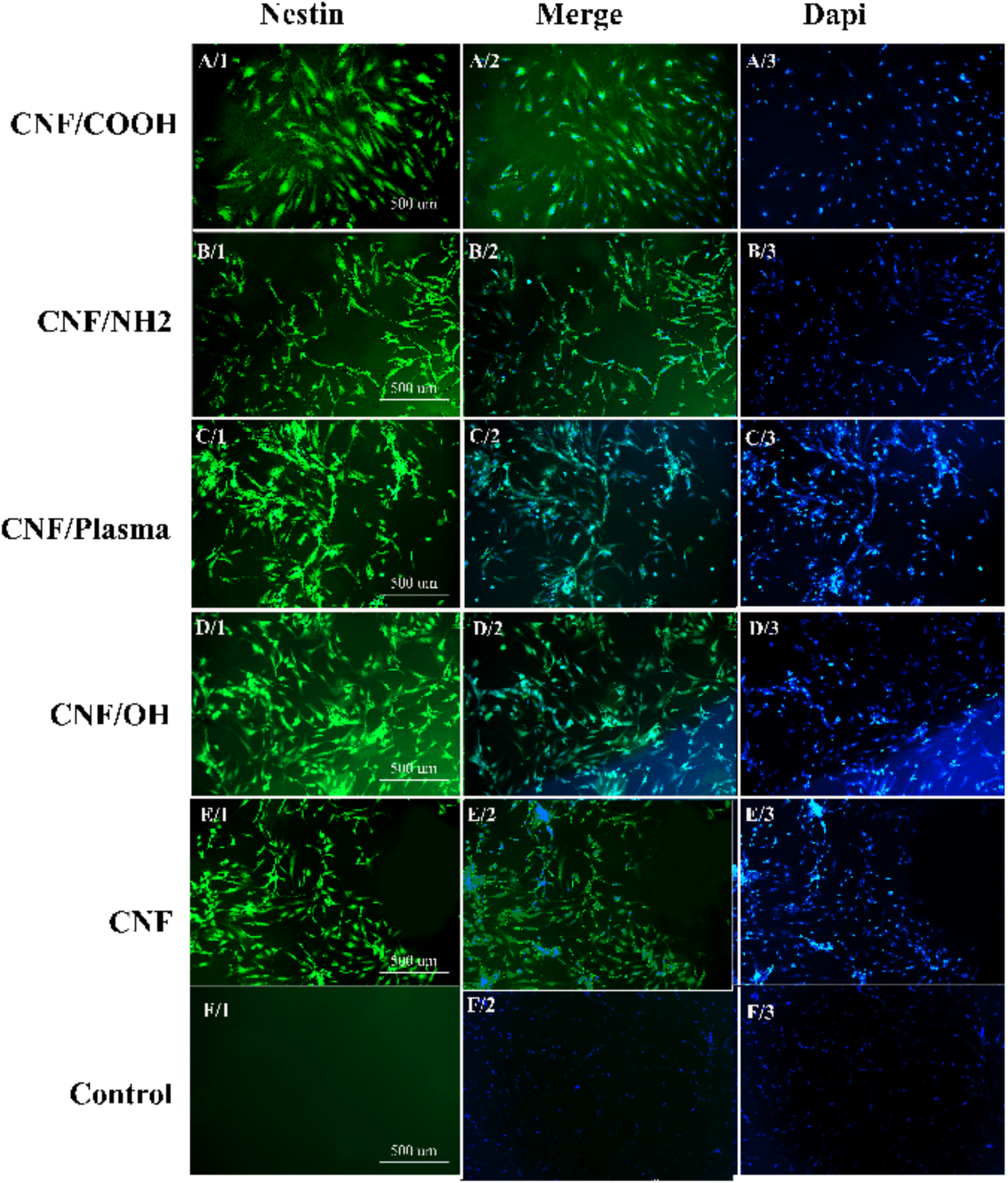
Immunofluorescence staining of nestin. (a, b, and c) CNFs-COOH, (d, e, and f) CNFs-NH_2_, (g, h, and i) CNFs-OH, (j, k, and l) CNFs-Plasma and (m, n, and o) CNFs. The scales are 100 μm.

#### 2.2.2. Electrical stimulation of hAMSCs through CNFs

The ES was conducted to stimulate differentiation of neural-like cells from the hAMSCs and the results were evaluated using real-time PCR (Figure 8) and immunofluorescence staining (Figure 9). The results showed that hydroxylated CNFs using oxygen Plasma resulted in higher genes (Map2, Tub B, Nestin, and GFAP) expression than the bare, aminated and Carboxilated CNFs. The differences for Tub B and GFAP were statistically significant (**: and ***:) while for Nestin and MAP2 were not significant. The higher effects of Plasma on the gene expression under the ES can be attributed to the more hydrophilicity of the resulting CNFs.

**Figure 8:**
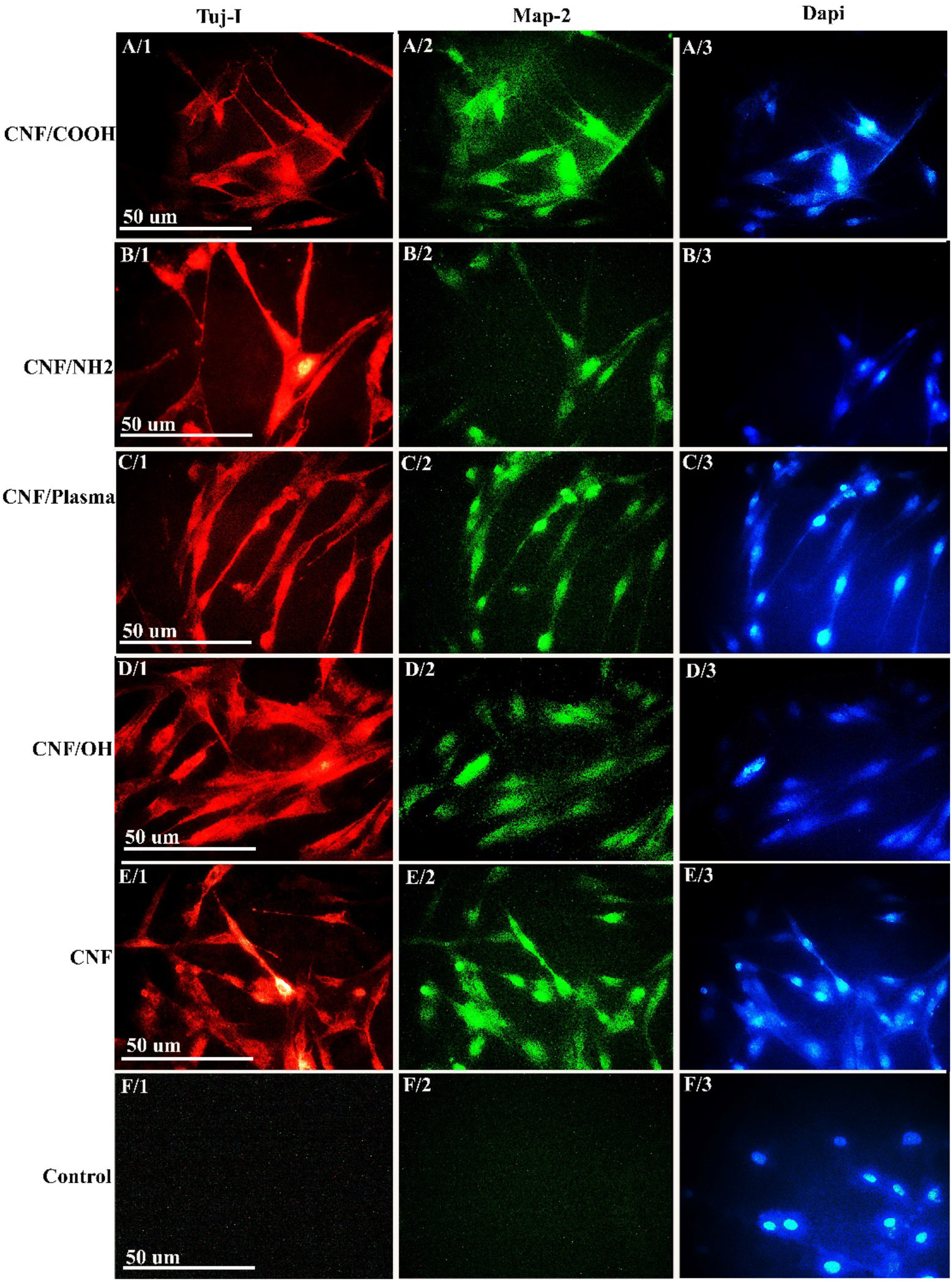
Immunofluorescence staining of Map2 and Tuj β. (a, b, and c) CNFs-COOH, (d, e, and f) CNFs-NH_2_, (g, h, and i) CNFs-Plasma, (j, k, and l) CNFs-OH and (m, n, and o) hAMSC as negative control. The scales are 100 μm.

The effect of the ES through the CNFs on neural protein expression was visualized using the immunofluorescence staining (**Figure 9**). It was observed that the ES through the treated CNFs with various surface functionality induced the expression of Nestin, Map2 and Tuj 1 proteins by the hAMSCs. These observations revealed that the ES induced neural proteins apart from the surface functionality of CNFs.

The results show that functionalization of the surface increases the percentage of differentiation, which can be due to increased cell interaction with the scaffold surface. The positive effect of plasma and amine groups on neural differentiation has been suggested, which can be due to better scaffold-cell interaction, which leads to improved and accelerated differentiation process. Plasma group shows the highest value, which can be due to further increase in surface hydrophilicity and the least structural changes of scaffold were observed with plasma-based method.

Adhesion, migration, and differentiation of cells depend to a great extent on the surrounding microenvironment, physical and chemical properties of the biological material surface. Hydrophilicity and chemical structure are essential factors affecting stem cell behavior. Studies have shown that mRNA expression, cell morphology, fibronectin and vitronectin levels, and MSC behavior change with tailoring the surface chemistry, thus affecting cell behavior and differentiation(25). Hydrophilic biomaterials cause absorbing more proteins and increase the transport and excretion of food materials(26, 27). Studies state cells in contact with the NH2 surface show a suitable cytoskeleton, high adhesion, and proliferation compared to glass plates (25). And by a higher density of amine on the surface increases cell connection, neuronal differentiation, and the formation of excitatory synapses. Compared to other chemically active groups, the amine group significantly promotes the adhesion and migration of NSCs due to affinity of the positive charge of *aminated* surface to the negative charge of membrane (28). In addition, studies have shown that a change in the carboxyl group’s density regulates cell adhesion and guides axonal growth through effects on the hydrophilicity of biological materials (29).

The XRD peak results show an increase in the width of the (002) band, which is associated with a decrease in the size of graphite crystals. In the CNF/plasma sample, there were fewer microstructural changes. The solvent type and time effect were effective on the surface structural changes. It seems that most of the changes in the amine group are caused by acid nitric solution (30).

In our studies, we observed that ID/IG ratio increases, so the ratio of boundaries-to-planes increases, which confirms the decrease in the size of graphite crystals and domains. These results are consistent with the XRD results. Of course, the structural changes are minimal, and the network structure has been preserved during the functionalization process.

By functionalizing nanofibers, electrical conductivity decreased. According to the XRD and Raman data, crystallinity reduction and structural defects in the graphite crystal and the formation of functional bonds with carbon, such as the C-O bond, which has replaced the C=C bond, can be mentioned as the reasons for reducing the conductivity of nanofibers (30).

Since the transfer of π electrons takes place in the form of sp2 hybridization, in the functionalized forms, it becomes a full sp3 bond, which decreases the speed of electron movement and conductivity(30).

These structures have recently been used in neurology discussions, but there are concerns about the toxicity of these structures, and they must be investigated.

Mirzai investigated the toxicity of carbon nanofibers on hEnSCs, and no toxicity was seen on the cells. The cells had a good proliferation on the scaffold, and the toxicity was below 10%, which can be used for biomedical applications(31).

In a study, they examined the compatibility of Schwann cells and neuroblasts on CNFs scaffolds under UV and plasma. They found that the survival of cells on plasma scaffolds is better than on UV and stated carbon nanofiber scaffolds. It is a suitable scaffold to support the connection and proliferation of nerve cells, and Schwann cells showed proper growth and proliferation on these scaffolds (32).

Some studies showed that scaffold conductivity is very important for nerve cells, and it has been reported that a carbon nanotube scaffold with conductivity less than 0.3 S/cm leads to neuronal growth. While not seen in higher conductivities (33).

In our study, there was no direct relationship between electrical conductivity and the degree of differentiation with electrical stimulation.

## 3. Conclusion

neural tissue engineering has emerged as a multi-disciplinary field combining material sciences, biomedicine, and engineering sciences/technologies to improve the conventional treatment modalities and eliminate their limitations. Neural tissue engineering requires sophisticated structure as the scaffold and potent biological, chemical, and/or physical cues to enhance the low regeneration potential of neural tissues. Electrical stimulation has shown promising results in the clinic to ameliorate neural damages. Electrical stimulation using the electroconductive scaffolds has fascinating outcomes due to localized and effective stimulation of the cells. Accordingly, in the current study, we fabricated CNFs as the scaffold for electrical stimulation of haMSCs to differentiate them toward neural cells. The surface of CNFs was treated using different chemicals method to introduce biologically favorable surface functional groups to improve the hydrophilicity of CNFs. The results showed that the oxygen plasma treatment has the highest effects on the hydrophilicity of the CNFs and induced the lowest adverse effect on the structure of CNFs. The aminated CNFs have shown the highest proliferative effects on hAMSCs in the absence of electrical stimulation. On the other hand, the Plasma treated CNFs exhibited the highest neural gene expression under electrical stimulation. This study showed that the electrospun CNFs have fascinating properties beneficial for neural tissue engineering and the surface of CNFs can be modified using a different method to be tailored for biomedical applications. The combination of CNFs with electrical stimulation can be considered a promising neural tissue engineering approach.

## 4. Materials and methods

### 4.1. Chemical and reagents

Poly(acrylonitrile) (PAN, MW: 150,000 gmol^−1^ was gifted by Polyacryl Company, Esfahan, Iran. Nitric acid 65%, *N,N*-dimethylformamide (DMF), sodium hydroxide (NaOH) and Ethylenediamine (EDA, ≥99 %) were purchased from Merck (Darmstadt, Germany). Dulbecco’s modified Eagle’s medium: nutrient mixture F-12 (DMEM/F12), fetal bovine serum (FBS), Pen-Strep(Penicillin-Streptomycin) and Trypsin-EDTA were Gibco (USA). 3-(4,5-dimethylthiazol-2-yl)-2,5-diphenyltetrazolium bromide (MTT) and Lactate dehydrogenase (LDH) assay kit (LDH Cytotoxicity Detection KitPLUS) were obtained from Roche (Germany). MG-63 cell line (National Cell Bank of Iran (NCBI), Pasteur Institute, Tehran, Iran),

### 4.2. CNFs fabrication

CNFs were obtained from electrospun PAN nanofibers after heat treatment. Briefly, PAN was dissolved in DMF with the concentration of 9 wt.% and stirred for 24 hr at 60 °C to obtain a clear and homogenous solution. The obtained solution was converted to nanofibers using the electrospinning device (Fanavaran Nano Meghyas Ltd., Co., Tehran, Iran) with the electrospinning parameters of applied voltage: 20 kV, feeding rate: 1 mL/hr, and nozzle-to-collector: 10 cm.

The fabricated PAN nanofibers were converted to CNFs using the heat treatment (34).

### 4.3. CNFs surface functionalization

Different methods were applied to introduce various surface functionality on the fabricated CNFs. First, the CNFs were treated with concentrated nitric acid (65 %) for 1, 2, and 4 hr at around 100 °C to introduce carboxylic groups on CNFs. The treated CNFs were removed from the acid, washed several times with deionized (DI) water, and dried at 70 °C. Second, amine functionalization was carried out using the treatment of CNFs with EDA. CNFs (200 mg) was treated with EDA (5 mL) for 40 min, then 2 mL DI water was added and incubated for 2 hr. Then, the aminated CNFs were washed with DI water and dried at 70 °C. Third, CNFs were treated with NaOH (5 mol) for 24 hr at 37 °C, then removed, washed with DI water, and dried at 70 °C. Fourth, oxygen plasma treatment was conducted on CNFs using a plasma instrument (Plasma CleanerPDC-32G-2(30 V)) with an oxygen flow of 50 mL/min for 30 and 60 s.

**Schematic 2:**
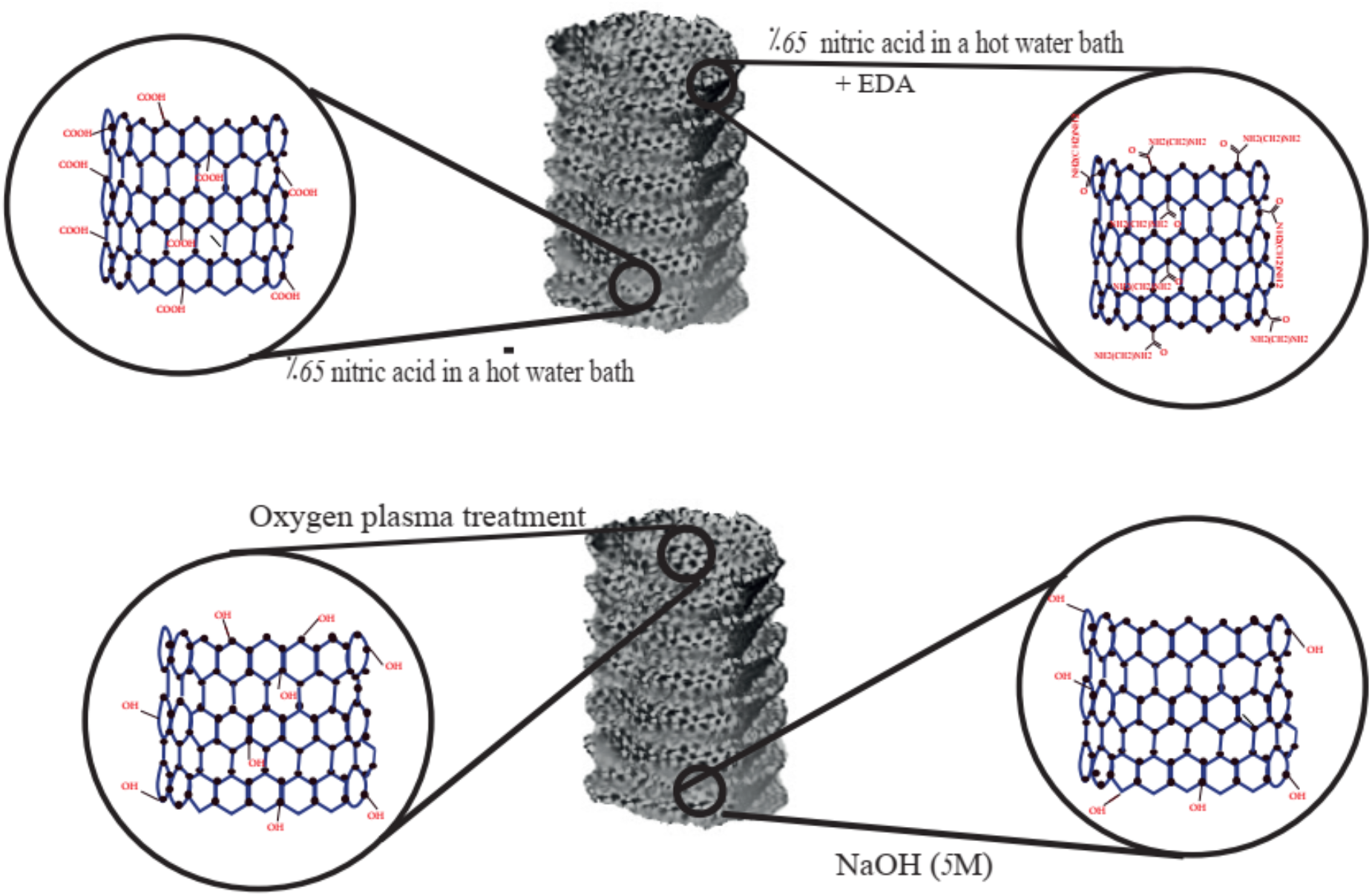
CNFs surface functionalization

### 4.4. Characterizations

The morphology of the CNFs was visualized using scanning electron microscopy (SEM, Philips XL 30, at 25.0 kV) after sputter-coting with gold using a sputter coater (SCD 004, Balzers, Germany). The CNFs diameter was measured using ImageJ software (NIH). Fourier transform infrared (FTIR) spectrum of the treated CNFs was obtained using Shimadzu 8101 M FTIR (Kyoto, Japan). Raman spectroscopy was conducted to assess the internal structure of CNFs using a Teksan Takram P50C0R10 Raman spectrometer with the laser power of 0.5–70 mW and the laser wavelength of 532 nm. The X-ray Diffraction (XRD) pattern of the CNFs was recorded using a Siemens D5000 diffractometer (Aubrey, TX) (Cu Kα radiation with λ = 1.5406 Å, 40 kV, 30 mA, 2θ range from 10° to 80° and the step size of 0.05°/s). A 4-point probe multi-meter (Signatone SYS-301) was used to measure the electrical conductivity of CNFs. The water contact angle (WCA) value of CNFs was obtained using a contact angle measuring instrument (Iran). The mechanical properties of CNFs were evaluated based on the tensile strength method according to ASTM-D 5024-95a standard using a Santam mechanical testing machine (Karaj, Iran) with a cross-head speed of 0.5 mm/min.

### 4.5. Biological studies

#### 4.5.1. Cell viability/proliferation and morphology on CNFs

The possible toxicity of the CNFs was evaluated using the in-direct MTT assay protocol and direct LDH assay protocol. For the in-direct MTT assay protocol, 100 mg of the sterilized CNFs (using ethanol 70% for 2 hr and UV radiation) with different surface treatments were incubated with 1mL FBS-free DMEM/F12 cell culture medium at 37 ° for 24, 28, and 72. The obtained extracts were incubated with pre-seeded hAMSCs for 24 hr and the possible toxicity was measured using the MTT assay kit. CNFs-free DMEM/F12 cell culture medium incubated with the same conductions was ued as the control. For the direct toxicity assessment, the CNFs were punched into disks, put on the bottom of a 96-well plate, and sterilized. A number of 5,000 cells/well in 100 μL DMEM/F12 cell culture medium containing FBS (1.0 %) and antibiotics was seeded on the CNFs and incubated for 24, 48, and 72 hr at 37 °C in a humidified cell incubator. Then, the cell culture mediums were transferred to another plate and the amount of LDH enzyme leaked from the cells under incubation with CNFs was measured using the LDH Activity Assay Kit as the direct toxicity indication (35). Using the same method, the total LDH content of grown cells on CNFs (after lysing with the buffer) was measured as the indication of cell proliferation (36). The morphology of the cells on CNFs after 24 cell-seeding was visualized using SEM after fixing, dehydrating, and sputter coating with gold.

#### 4.5.2. Electrical stimulation of hAMSCs through CNFs

The electrical stimulation of hAMSCs-cultured on the CNFs was conducted using a homemade electrical stimulation setup customized for a 24-well cell culture plate connected to a Victor 2015H Signal/Function Generator (Schematic 1). The CNFs were put at the bottom of wells and sterilized. A number of 20,000 cells/well was seeded on the CNFs and incubated for 48 hr to completely attach to CNFs. Then, the electrical stimulation was carried out for 7 days 10 min each day by a 1.5 mA current with a frequency of 500 Hz and CMOS waveform. The effects of the electrical stimulation on the neural differentiation of hAMSCs were investigated using real-time PCR and immunofluorescence staining. Using the real-time PCR the expression of Glial fibrillary acidic protein (GFAP), Microtubule Associated Protein2 (Map2), Nestin, and Neuron-specific class III beta-tubulin (Tuj 1) genes were evaluated and the immunofluorescence staining was conducted to assess the expression of Nestin on the cells.

### 4.6. Statistical analysis

The experiments were conducted in triplicate and the obtained data were reported as mean ± standard deviation (SD). The statistical analysis was carried out using SPSS software version 10.0 and the one-way analysis of variance (ANOVA) was conducted as the posteriori test.

## Acknowldment

We thank our colleagues in Tehran University of Medical Sciences and National Cell Bank of Iran, Pasteur Institute of Iran. This work was supported by Tehran University of Medical Sciences, grant no. 98-01-87-41015.

## Conflict of intrest

The authors declar they have no conflict of intrest.

